# The Mexican Cavefish Mount a Rapid and Sustained Regenerative Response Following Skeletal Muscle Injury

**DOI:** 10.1101/2023.02.05.527207

**Authors:** Luke Olsen, Huzaifa Hassan, Sarah Keaton, Nicolas Rohner

## Abstract

Physical injury and tissue damage is prevalent throughout the animal kingdom, with the ability to quickly and efficiently regenerate providing a selective advantage. The skeletal muscle possesses a uniquely large regenerative capacity within most vertebrates, and has thus become an important model for investigating cellular processes underpinning tissue regeneration. Following damage, the skeletal muscle mounts a complex regenerative cascade centered around dedicated muscle stem cells termed satellite cells. In non-injured muscle, satellite cells remain in a quiescent state, expressing the canonical marker *Pax7* (Chen et al. 2020). However, following injury, satellite cells exit quiescence, enter the cell cycle to initiate proliferation, asymmetrically divide, and in many cases terminally differentiate into myoblasts, ultimately fusing with surrounding myoblasts and pre-existing muscle fibers to resolve the regenerative process (Chen et al. 2020).

---

The innate immune system, specifically those of the myeloid lineage, are crucial for the sequential steps described above (Tidball et al. 2017). For example, within the first few hours of muscle damage, neutrophils and macrophages infiltrate the damaged skeletal muscle to mediate the repair process; including phagocytosis of cellular debris, secretion of cytokines regulating satellite cell dynamics, and temporal control of the pro- and anti-inflammatory phases (Tidball et al. 2010). However, investment into the innate immune system is energetically costly resulting in many species decreasing investment in the innate immune system under nutrient-limited environments (McDade et al. 2016). Whether this reduced investment into the innate immune system results in a decreased capacity to mount a regenerative response following tissue damage remains unclear. To this point, we utilized the emerging evolutionary model, *Astyanax mexicanus*, to investigate the consequence of shifts in immune system investment on skeletal muscle regeneration.

The *Astyanax mexicanus* is a single species, comprised of river-dwelling surface fish and cave-dwelling cavefish. ∼160,000 years ago, the ancestral surface fish colonized the surrounding caves resulting in multiple independently derived cave populations (Herman et al. 2018). These caves are completely devoid of light, resulting in diminished primary producers and a subsequent general lack in biodiversity. These different selective pressures have led to many cave-specific morphological and physiological adaptations (Krishnan et al. 2020; Stockdale et al. 2028; Peuß et al. 2020; Olsen et al. 2022). For example, we found that the diminished macroparasite diversity in certain caves affected the evolution of the immune investment strategy and the sensitivity of the immune system towards immunological stimuli (e.g. lipopolysaccharides; LPS) of cavefish (Peuß et al. 2020). Here, the hematopoietic niche of cavefish consists of less innate immune cells (e.g., neutrophils and monocytes) than the surface fish and we found that this reduction of proinflammatory cells is compensated by a prolonged pro-inflammatory response through stronger and more sustained expression of proinflammatory cytokines such as *il6, tnfα, il1β*, and *g-csf* upon exposure to LPS (Peuß et al. 2020). Similarly, injuring the heart of *A. mexicanus* resulted in an elevated immune response within cavefish relative to surface fish – a phenomenon thought to underlie their inability to fully regenerate their cardiac tissue (Stockdale et al. 2018). To this point, we sought to address whether a similar decrease in regenerative potential persists within the cavefish skeletal muscle following injury via analyzing the transcriptional responses after cardiotoxin injection in both surface fish and cavefish. The fact that cavefish and surface fish populations are of the same species makes comparative transcriptomic approaches robust and informative (Krishnan et al. 2020).

## Methods

### Cardiotoxin injection and muscle collection

All fish used were adult (1-2 years old) and reared at similar tank densities. Laboratory-reared surface fish and cavefish (of the “Pachón” population) were fasted overnight and anesthetized in ms222 for ∼30 seconds or until movement stopped. Following anesthetization, fish were injected with 0.3 mg/ml cardiotoxin dissolved in 1xPBS (pH 7.4). Control fish were similarly anesthetized and injected with 1xPBS (pH 7.4). Injections were made immediately anterior to the dorsal fin to decrease the likelihood of impacting swimming behavior. No obvious changes in swimming behavior were observed. Following injection, fish were transferred to their tank water to regain consciousness. Fish were then euthanized at 1 day post injury (dpi), 7dpi, and 14 dpi and skeletal muscle was immediately dissected and frozen in liquid nitrogen. The timepoints utilized within this study were determined based off previous literature and histological analysis of surface fish skeletal muscle regeneration (Figure S2). Surface fish are considered as the ‘control’ group, and we thus sought to determine the regenerative process first within surface fish and identify timepoints which best reflect the central components of skeletal muscle regeneration (i.e. immune cell infiltration and new muscle fiber formation). We determined timepoints ranging from 1dpi- to-14dpi contained the most pertinent regenerative information and we thus used these timepoints for our follow-up muscle collection from a separate cohort of surface fish and cavefish for transcriptome analysis. All experiments and fish husbandry are approved under IACUC protocol 2021-129.

### RNA extraction and processing for RNA-sequencing

∼100mg of frozen tissue was homogenized in 1mL Trizol (Ambion) with triple pure M-Bio grade high impact zirconium beads in a Beadbug 6 microtube bead beater. RNA was extracted using standard phenol/chloroform extraction. The RNA pellet was cleaned with the RNeasy Mini Kit (Qiagen #74104) with on-column DNAse digestion (Qiagen #79256). Libraries were prepared according to the manufacturer’s instructions using the TruSeq Stranded mRNA Prep Kit (Illumina #20020594). The resulting libraries were quantified using a Bioanalyzer (Agilent Technologies) and Qubit fluorometer (Life Technologies). Libraries were normalized, pooled, multiplexed, and sequenced on an Illumina NextSeq-75-HO instrument as v2 Chemistry High Output 75bp single read runs. Following sequencing, Raw reads were demultiplexed into Fastq format allowing up to one mismatch using Illumina bcl2fastq2 (v2.20). Reads were aligned to the *Astyanax mexicanus* reference genome from University of California Santa Cruz with STAR aligner (v 2.7.3a), using Ensembl 102 gene models. Transcripts per million (TPM) values were generated using RSEM (v 1.3.0). Pairwise differential expression analysis was performed using Bioconductor package edgeR (v3.24.3 with R v3.5.2). Only protein coding genes and long non-coding RNAs (lncRNAs) were considered from the Ensembl 102 annotation. Only genes with counts per million expression ≥ 0.5 in at least 2 samples were kept for further analyses. Statistical significance was determined by fold change cutoff of 2 and false discovery rate (FDR) cutoff of 0.05 with the p.adjust function in R with the default choice Benjamini & Hochberg correction. Spearman correlations between all samples were determined and clustered. The correlation plot is based on the log2 transformed normalized gene expression values (TPM) that are generated using RSEM. The hierarchical clustering heatmap includes genes expressed above 2 TPM in at least one sample and is clustered based on Euclidean distance between samples. Gene Ontology (GO) Enrichment analysis was performed on differentially expressed genes identified in edgeR using R package ‘clusterProfile’.

## Results

### Global gene expression response to skeletal muscle injury

To characterize the regenerative response of *A. mexicanus*, we injected fish skeletal muscle with the necrotic agent cardiotoxin – shown to result in rapid and dramatic muscle necrosis (Seger et al. 2011 and Figure S2). Following injection, we collected skeletal muscle 1-day, 7-days, and 14-days post injury (dpi) followed by genome-wide analysis via bulk RNA-sequencing (Fig. 1A). In sum, we found cardiotoxin injection resulted in a more robust gene expression response at both the 1dpi, 7dpi, and 14dpi timepoints within cavefish relative to surface fish. Specifically, we identified 2,936 differentially expressed genes (DEG’s) at 1dpi within cavefish which markedly decreased to 944 DEG’s at 7dpi, and 1,656 DEG’s at 14dpi. Surface fish showed a similar, albeit reduced, increase in DEG’s at the 1dpi timepoint with a total of 1,851 DEG’s identified. This number decreased at 7dpi to 673 DEG’s, and continued to decrease at 14dpi with no DEG’s detected. In support of this, hierarchical clustering revealed the 1dpi samples clustered more distinctly than all other timepoints (Fig. 1B). Taken together, we find that cardiotoxin injection results in a robust change in gene expression within both cavefish and surface fish skeletal muscle, with a more extreme and sustained change in gene expression within cavefish.

**Figure 1.**
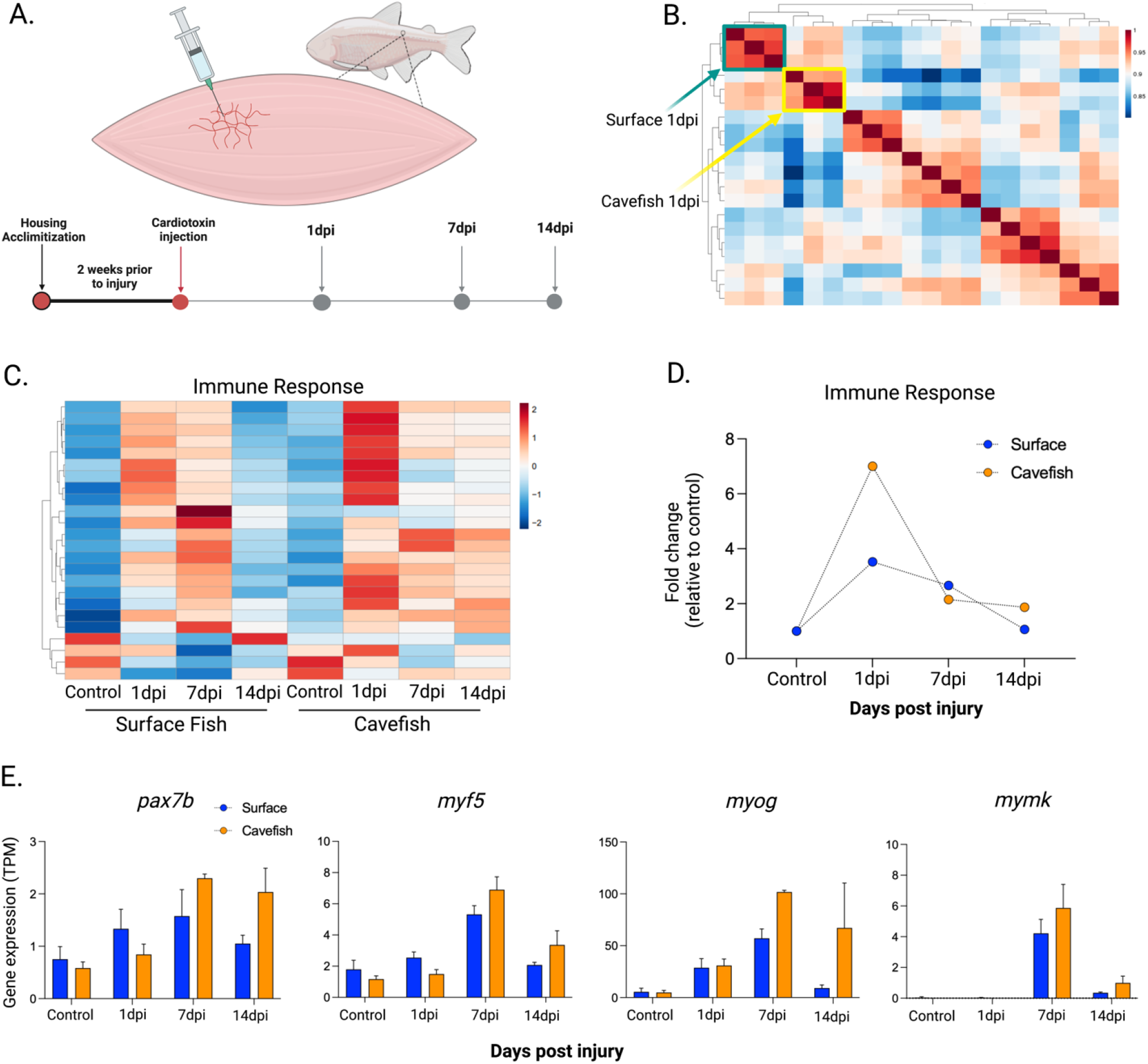
(A) Schematic of cardiotoxin injection and tissue collection at 1 day post injury (dpi), 7dpi, and 14dpi. (B) Hierarchical clustering heatmap of each sample analyzed with specific emphasis placed on 1dpi in both surface fish (green) and cavefish (yellow). (C) Heatmap of the genes identified within the immune response pathway of surface fish at the 7dpi timepoint and the corresponding genes within cavefish. (D) Average fold change of the genes identified within the immune response pathway (from Fig. 1C). (E) Relative gene expression of canonical satellite cell markers in transcripts per million (TPM).

### Immune response following skeletal muscle injury

Because of our recent findings that cavefish have decreased investment in the innate branch of the immune system – an essential component of muscle regeneration – we sought to explore gene expression changes underlying pro- and anti-inflammatory signaling. We first conducted a Gene Ontology enrichment analysis of all DEG’s at each timepoint and, as expected, identified multiple pathways enriched for the immune system. Specifically, we found “immune response” (*p*=0.0014), “adaptive immune response” (*p*=0.0016), and “positive regulation of immune response” (*p*=0.006) increased at 7dpi within surface fish (Fig. S1A). Cavefish showed a similar increase in immune-related pathways, such as “immune response” (*p*=0.0116) and “defense response” (*p*=0.0116), albeit at the 1dpi timepoint (Fig. S1B). As shown in Figure 1C, most genes within these pathways increased following injection, most dramatically at 1dpi and 7dpi, and tended to decrease back to baseline at 14dpi, though to a lesser degree in cavefish, a possible reflection of prolonged immune signaling following muscle damage. In fact, when pooling all genes identified within the “immune response” pathway, cavefish showed a greater increase in expression relative to surface fish (Fig. 1D).

In addition to those within the “immune response” pathway, we sought to characterize classical markers regulating inflammation previously shown to be differentially regulated between cavefish and surface fish, specifically the cytokines *il6, tnfα, il1β*, and *g-csf* (*gcsfa*). In agreement with our previous findings, cavefish had reduced basal expression relative to surface fish. Interestingly however, cavefish showed a robust increase in expression at 1dpi, having an ∼19-fold, 37-fold, 9-fold, and 6-fold increase in expression of *il6, tnfα, il1β*, and *g-csf*, respectively, relative to an ∼4-fold, 5-fold, 2-fold, and 1.1-fold increase, respectively, within surface fish (Fig. S1C). These data support our previous findings of cavefish having fewer innate immune cells (indicated by decreased expression at baseline), however, having heightened sensitivity following an inflammatory stimulus relative to surface fish – at least within the local site of injection (Peuß et al. 2020).

### Cavefish have an increased satellite cell response to injury

The ability to efficiently coordinate the inflammatory response following muscle injury is critical for functional satellite cell dynamics and the resolution of regeneration. Because cavefish appear to possess a more robust inflammatory response relative to surface fish following cardiotoxin injection, we sought to determine how this influences satellite cell dynamics, specifically the central genes regulating satellite cell quiescence (*pax7a/*b), activation/proliferation (*myf5/myod*), differentiation (*myod/myog*), and ultimate myoblast fusion (*mymk*) (Chen et al. 2020; Millay et al. 2013). As expected, many of these genes were increased in expression within both surface fish and cavefish following cardiotoxin injection. Surprisingly though, cavefish showed a more robust and sustained increase in expression relative to surface fish. For example, cavefish had a significant increase in *pax7b* at the 7dpi and 14dpi timepoint whereas surface fish increased but did not reach statistical significance (Fig. 1E). Additionally, cavefish increased expression of *myf5* (at 7 and 14dpi) and *myog* (at 1, 7, and 14dpi) following injury whereas surface fish only increased *myog* at 7dpi (Fig. 1E). Moreover, *mymk* showed a robust increase at the 7dpi and 14dpi timepoints within cavefish while increasing only at the 7dpi timepoint within surface fish. As such, in agreement with the immune system dataset, cavefish showed a more robust and sustained increase in genes orchestrating satellite cell dynamics following injury, a possible reflection of their heightened sensitivity to an external stimulus.

## Discussion

Here we provide evidence suggesting cavefish skeletal muscle initiates a ‘hyper-sensitive’ inflammatory response following local muscle injury. We reasoned this may result in dysregulation of the satellite cell response – similar to what is seen following cardiac injury within cavefish – but, in contrast to our expectations, cavefish demonstrated a more robust increase of markers regulating satellite cell proliferation, differentiation, and fusion when compared to surface fish. This may reflect cavefish being more sensitive to an injury stimulus and thus requiring a greater reliance on their satellite cell pool as compared to surface fish. While these data provide the necessary first step in delineating cavefish skeletal muscle regeneration, future work is required to confirm that our observations at the level of gene expression are in fact consequential toward muscle regeneration. For example, the sustained immune response seen within the cardiac tissue of cavefish results in scarring and fibrosis, a possible result within cavefish skeletal muscle. As such, histological analysis of cavefish and surface fish skeletal muscle at multiple timepoints following injury is required to determine whether our findings indicate a physiological or pathological response to muscle injury.

## Acknowledgements

We would like to thank the cavefish facility at the Stowers Institute for cavefish husbandry support, specifically Molly Miller, Elizabeth Fritz, David Jewell, Andrew Ingalls, Zachary Zakibe, Cole Biesemeyer, Hailey Gregory, and Diana Baumann. We thank Mark Miller for assistance with illustrations. We thank all core support provided by the Stowers Institute; specifically, Rhonda Egidy and Amanda Lawlor of Sequencing and Discovery Genomics. We would like to thank Robert Peuß for the critical review of our manuscript. NR is supported by institutional funding, NIH Grant 1DP2AG071466-01, NIH Grant R01 GM127872, NSF IOS-1933428, and NSF EDGE award 1923372.

---

**Figure S1.**
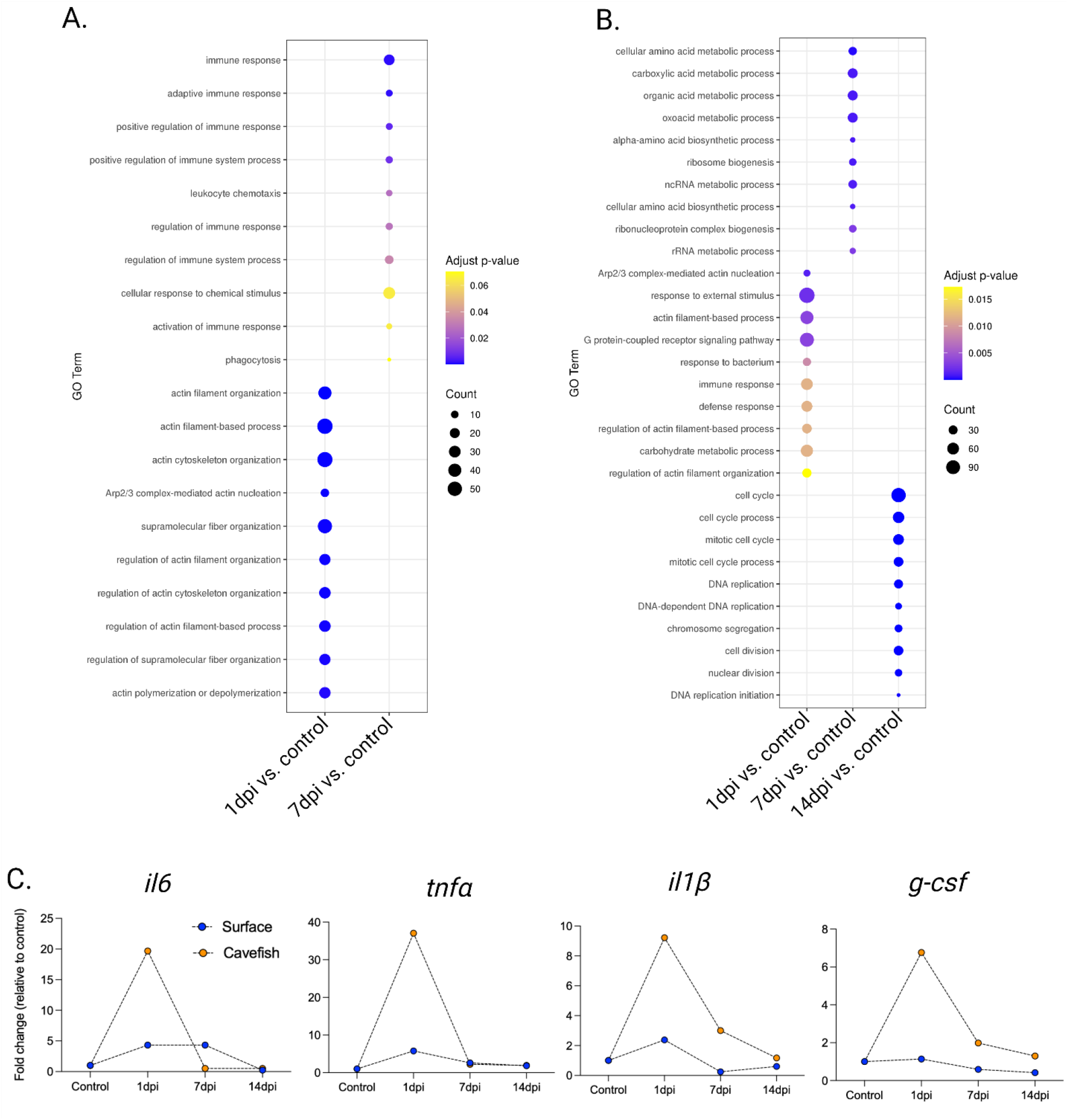
Gene Ontology enrichment analysis of the differentially expressed genes from 1dpi, 7dpi, and 14dpi relative to control (not injected) from (A) surface fish and (B) cavefish. Surface fish (Fig. S1A) did not have any DEG’s at the 14dpi timepoint relative to control. (C) Fold change of the pro-inflammatory cytokines *il6, tnfα, il1β*, and *g-csf*. Each timepoint is the mean of two-to-three biological replicates.

**Figure S2.**
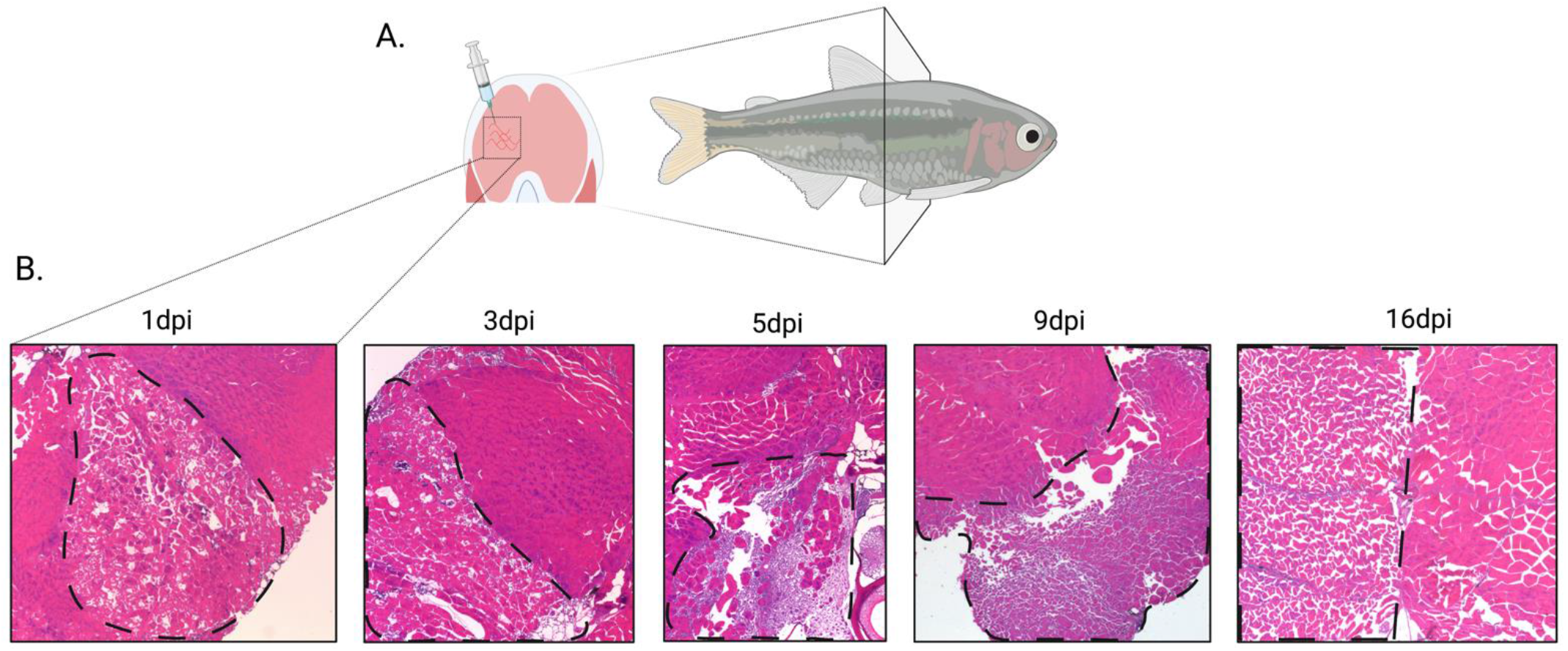
(A) Schematic of the site of skeletal muscle injection. (B) Representative images of muscle collected from the injured tissue at 1 day post injury (dpi), 3dpi, 5dpi, 9dpi, and 16dpi to determine the timepoint for the transcriptome analysis. The site of injury is demarcated by the black dashed line with the non-injured muscle directly adjacent. As mentioned within the Methods section, surface fish served as the control, and we thus determined the necessary timepoints for muscle collection of both cavefish and surface fish following histological analysis from surface fish at multiple timepoints post injury (shown above). As such, all images are from surface fish.

## Notes

### Competing Interest Statement

The authors have declared no competing interest.

